# uniForest: an unsupervised machine learning technique to detect outliers and restrict variance in microbiome studies

**DOI:** 10.1101/2021.05.17.444491

**Authors:** R.J. Leigh, R.A. Murphy, F. Walsh

## Abstract

Isolation Forests is an unsupervised machine learning technique for detecting outliers in continuous datasets that does not require an underlying equivariant or Gaussian distribution and is suitable for use on small datasets. While this procedure is widely used across quantitative fields, to our knowledge, this is the first attempt to solely assess its use for microbiome datasets. Here we present uniForest, an interactive Python notebook (which can be run from any desktop computer using the Google Colaboratory web service) for the processing of microbiome outliers. We used uniForest to apply Isolation Forests to the Healthy Human Microbiome project dataset and imputed outliers with the mean of the remaining inliers to maintain sample size and assessed its prowess in variance reduction in both community structure and derived ecological statistics (α-diversity). We also assessed its functionality in anatomical site differentiation (pre- and postprocessing) using principal component analysis, dissimilarity matrices, and ANOSIM. We observed a minimum variance reduction of 81.17% across the entire dataset and in alpha diversity at the Phylum level. Application of Isolation Forests also separated the dataset to an extremely high specificity, reducing variance within taxa samples by a minimum of 81.33%.

It is evident that Isolation Forests are a potent tool in restricting the effect of variance in microbiome analysis and has potential for broad application in studies where high levels of microbiome variance is expected. This software allows for clean analyses of otherwise noisy datasets.

## Introduction

Due to their sensitivity to environmental conditions, microbiomes are often highly variable between studies, between samples of the same study, and within individuals at different timepoints (*eg*. Falony *et al*., 2016). Statistical and bioinformatic tests to detect suitable differentiable markers are continuously innovated, however microbiome studies are still impacted by the presence of highly variant outliers (Hahn and Zemanick, 2019). While considerable effort has been invested in outlier processing in human samples (Montassier *et al*., 2018), to our knowledge, a robust, phylogeny independent outlier detection and processing method has not been developed that can be universally applied to microbiome data. Microbiome phylogenetic metrics such as UniFrac (Lozupone *et al*., 2006) can be immensely informative, however due to the difficulties in accurately resolving phylogenies, the ever-changing reclassification of microbial taxonomy, and the difficulty in classifying unknown taxa observed in samples, researchers may be reticent to use these metrics (Rappé and Giovannoni, 2003; Liao *et al*., 2020). Outlier removal is a controversial topic in data science, and it is often common practice not to remove outliers post detection, however, as extreme variance in an observed microbiome is likely to indicate dysbiosis or due to the influence of an environmental factor that was not intended to be assessed (Falony *et al*., 2016), their removal, if desired, is legitimate. To overcome the hurdle of outliers, we assessed the proficiency of Isolation Forests (Liu *et al*., 2008) to reduce the impact of outliers. This article also assesses the effect of different simple imputation methods on datasets post outlier removal. The Python script used to generate this data uses the Scikit-Learn library (Pedregosa *et al*., 2011) and shall be provided as an iPython notebook so any user can utilise this tool with Google Colaboratory (https://colab.research.google.com/) regardless of their coding experience.

A detailed manual describing each module and how to run the software is available at the code repository (https://github.com/RobLeighBioinformatics/MicrobiomeOutliers).

## Methods

### Assumptions

This procedure is an **unsupervised** machine learning method and so is suitable for detecting outliers from **similar samples** (*eg*. from the same timepoint at the same location) so that extreme outliers between data are not dominant.

### Data scaling (normalisation)

Sequence data often displays disparate sequencing depths between samples. As such, data scaling (normalisation) is employed to remove such disparity and allow for more representative data comparison. The goal of this procedure is to ensure that the sum of all taxa in each sample sum to the same value. Each taxon in microbiome samples in a given study can easily be normalised using their 16S counts (or other taxonomic identifiers) using the formulae:

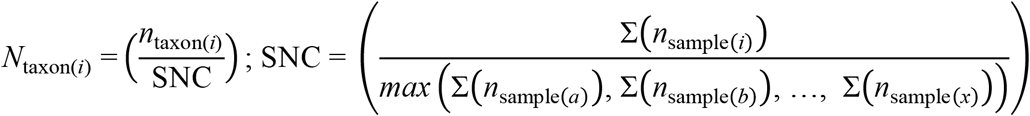

Where:

*N*_taxon_(*i*): Normalised reads for taxon *i*

*n*_taxon_(*i*): Observed reads for taxon *i*

∑(*n*_sample_(*i*)): Sum of all observed reads in sample *i*

SNC: Sample normalisation constant

This normalisation procedure serves to reduce the potential variance and standard error between datasets to yield a more representative sample for comparison.

### Outlier processing

There are several procedures to detect and process outliers, for example One Class Support Vector Machines (Dumais *et al*., 1998), Elliptic Envelope (Rousseeuw and Van Driessen, 1999), Local Outlier Factor (Breunig *et al*., 2000), and Isolation Forest (Liu *et al*., 2008). For the purposes of this manuscript, we are focussing on Isolation Forest as it is relatively fast, does not require data to follow a Gaussian distribution, and handles small datasets highly effectively. The strengths of Isolation Forest resound in its efficiency in *global* outlier detection (outliers from the entire group; Figure 1). Comparatively, Isolation Forest does not effectively distinguish *local* outliers (outliers within subgroups in a given group). This distinction should not have a negative impact on microbiome studies as this process is intended to be applied to similar samples (groups) only prior to *post hoc* comparisons.

**Figure 1:**
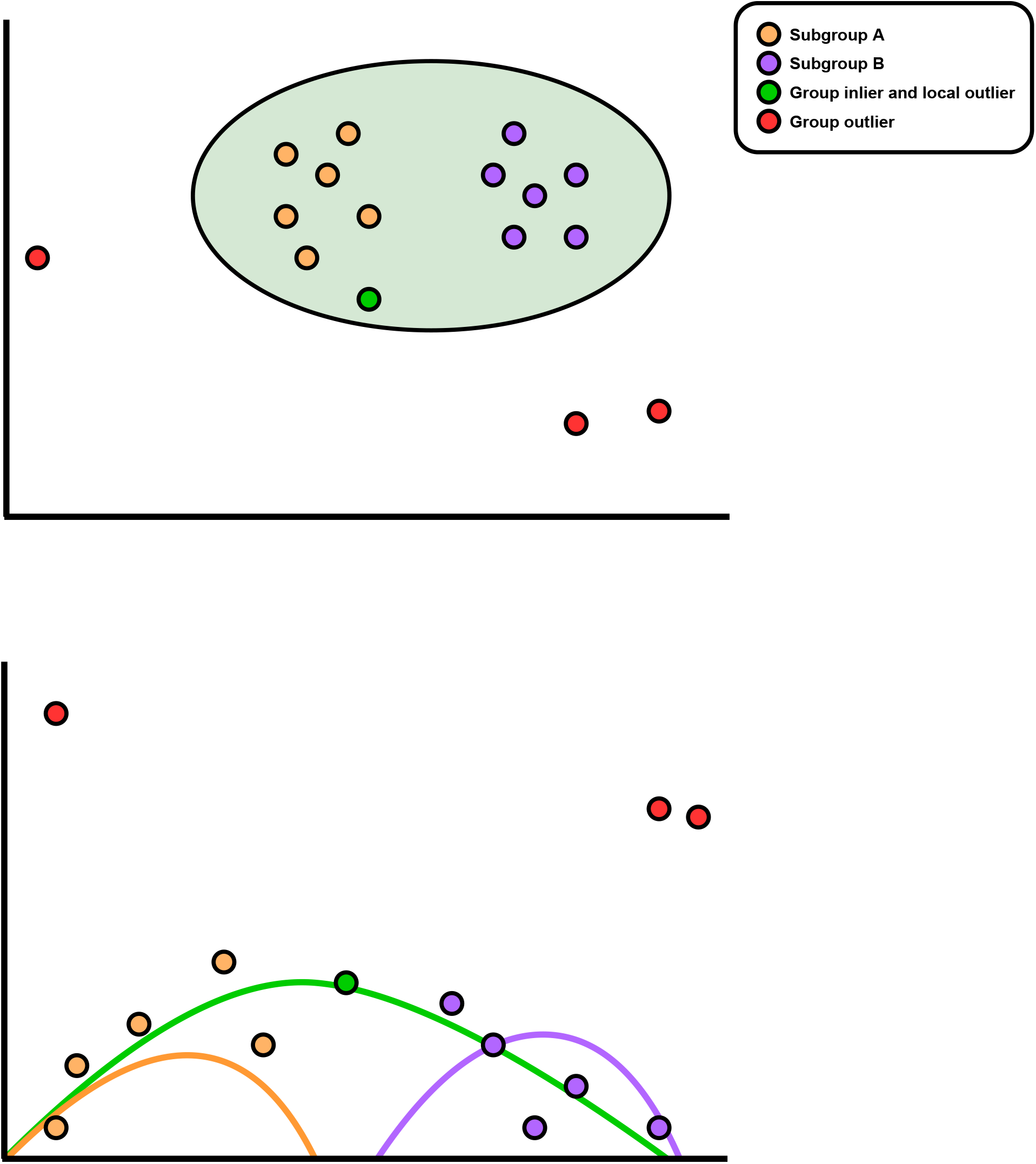
Difference between group outliers and local outliers. The upper portion of the graph illustrates how a group (pale green) can be locally subcategorised into local subgroups (pale orange and pale purple). The dark green sample represents a member of the group that could not be sub categorised further rendering it a group inlier but local outlier. Red samples are group outliers. The lower portion of the graph represents the upper portion in linear format.

### Isolation Forest

Unlike many other (supervised) outlier removal strategies which operate by constructing a profile of “normal data”, Isolation Forest does not rely on training data (instead working with only input samples). As this procedure avoids computationally expensive training steps with potentially biased data, Isolation Forest provides an accurate, rapid, computationally non-expensive strategy for outlier processing.

### Application to Healthy Human Microbiomes

#### Dataset construction

The 16S V3-V5 region dataset of Healthy Human Microbiome project (Huttenhower *et al*., 2012) was downloaded from MicrobiomeDB (Oliveira *et al*., 2018). All non-bacterial reads were removed and *m* × *n* matrix was constricted by imputing all missing data with 0 and aggregating all reads within each respective Phylum. Each sample was annotated with its sampling site (as a header row) obtained from the original publication: ‘Colon’, ‘External_ear’, ‘External_naris’, ‘Gingiva’, ‘Hard_palate’, ‘Median_vaginal_canal’, ‘Mouth_mucosa’, ‘Oral_opening’, ‘Palatine_tonsil’, ‘Posterior_fornix_of_vagina’, ‘Skin_of_elbow’, ‘Throat’, ‘Tongue’, ‘Vagina_orifice’ or ‘Calcareous_tooth’. The ‘Calcareous_tooth’ category was further annotated as ‘Subgingival_dental_plaque’ or ‘Supragingival_dental_plaque’ yielding 16 total groups. This procedure was conducted at the Phylum level so as to provide the most stringent example as it was expected that groups would be most similar at higher levels.

#### Parameter testing

The dataset was processed using uniForest to test the performance of each impution parameter (arithmetic mean, geometric mean, harmonic mean, and median) with iForest bootstrapping and data scaling (*n*_tests_ = 16). Data impution is used to replace outliers with a representative value so as to keep sample sizes constant pre- and postprocessing.

The performance of a given impution metric was assessed using one of two processes. The first process focusses on reducing “contamination” by increasing distance (dissimilarity) between groups.A Bray-Curtis dissimilarity matrix (Bray and Curtis, 1957) was constructed between each pair of groups (*n*_groups_ = 16) yielding 120 dissimilarity matrices (_*n*_*C*_*r*_ = _16_*C*_2_ = 120). Each matrix was transformed to a dissimilarity score (*D*) using ANOSIM (Clarke, 1993), where scores approaching -1 indicated greater group similarity and scores approaching 1 indicated greater group dissimilarity. Normalised differences (*D*’) between the raw data and processed dissimilarity scores (*D*_raw_ *vs. D*_processed_) using the formula:

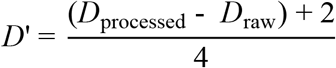

As ANOSIM scores fall between {-1, 1}, differences can also only fall between {-2, 2}, by adding maximum possible score (2) to the difference and dividing by the range (4), *D’* falls between {0, 1} where a *D’* < 0.5 indicates a decrease in performance, a *D’* = 0.5 indicates no change in performance, and a *D’* > 0.5 indicates an increase in performance (Figure 2). This score accounts for all possible scenarios and thus may appear to be quite punitive. For example, a *D*^’^ = 1 is only achievable in a scenario where a *D*_raw_ = -1 (all samples within a group are completely eclipsed by samples within another group) and a *D*_processed_ = 1 (no sample overlap between groups) are initially observed (Figure 2). To quantify the magnitude of this score a Kruskal-Wallis *H* test (Kruskal and Wallis, 1952) is performed between both groups (H_0_:η_(*a*)_=η_(*b*)_:H_A_:η_(*a*)_≠η_(*b*)_). A Bonferroni-Dunn correction was applied (*n* = 16), and a *P*_BD_ ≤ 0.005 was considered statistically significant. The Kruskal-Wallis *H* test was chosen over other statistical comparisons as it does not require data to follow a Gaussian distribution or equivariance between groups and is powerful even when using small sample sizes (*n* ≥ 5) thus allowing for more reliable comparability between groups (Table 1).

**Figure 2.**
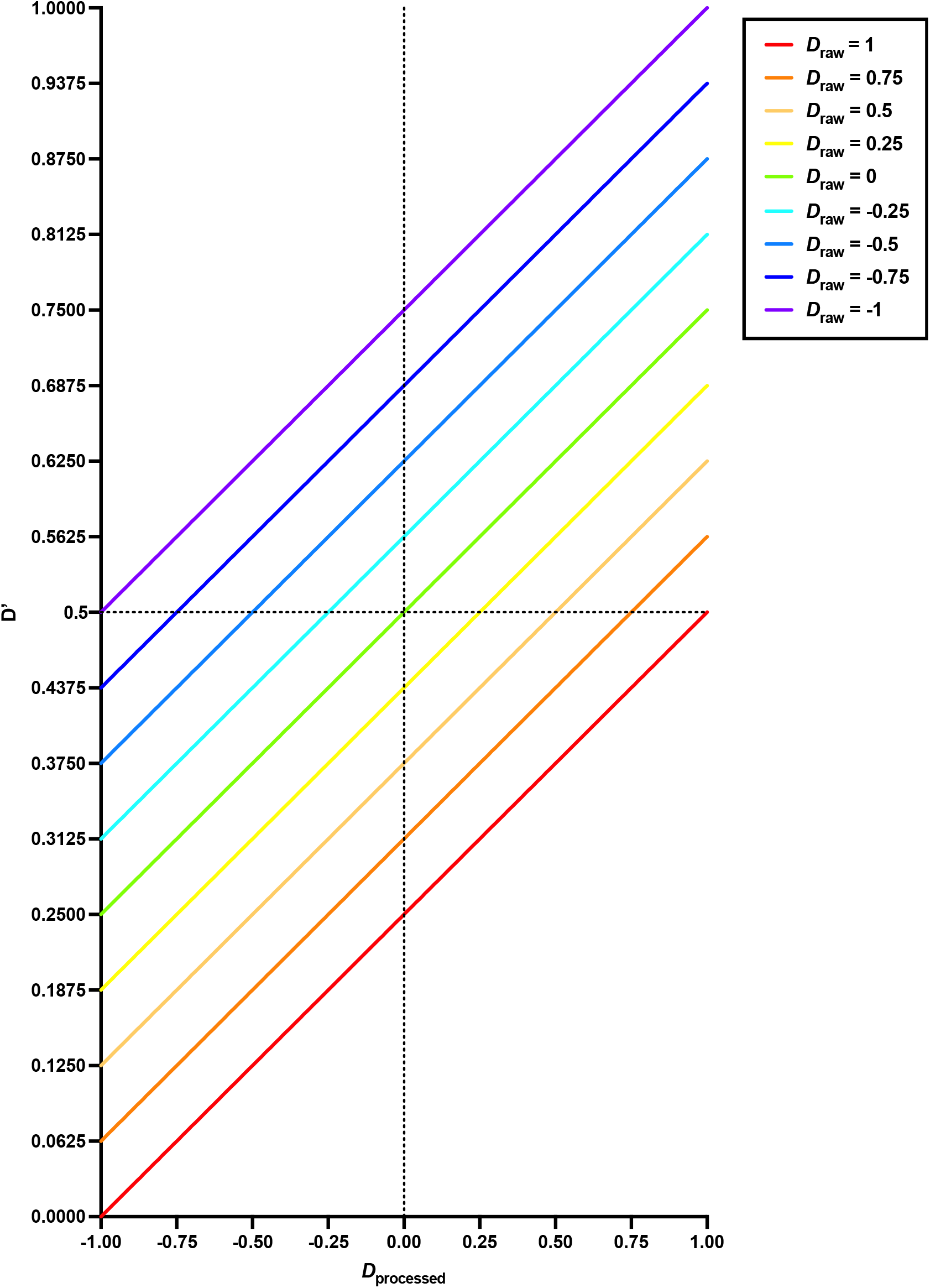
Impution metric performance scores. The horizontal dashed line illustrates the point at which an improvement between *D*_raw_ and *D*_processed_ is achieved. The vertical line illustrates where at 50% separation between groups in the processed dataset is achieved.

The second performance process focusses on the reduction of distance between samples in the same group and the reduction of variance in distances between samples. Separate principal component analyses (PCA) were performed for each group on the raw and processed datasets. The three most prominent components were taken as *x, y*, and *z* Cartesian coordinates for each sample and weighted by multiplying each coordinate by its explained ratio. Distances (*d*) between each weighted sample were computed using the formula:

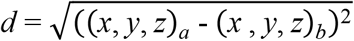

Distance reductions were assessed by comparing the distribution of distances between the raw dataset and processed dataset using a Kruskal-Wallis *H*-test (Table). A Bonferroni-Dunn correction was applied (*n* = 16), and a *P*_BD_ ≤ 0.005 was considered statistically significant. It has been previously recommended that α ≤ 0.005 is preferable to α ≤ 0.05 to ensure greater statistical power and accuracy (Benjamin *et al*., 2018).

Treatment effect (comparison of variation) was assessed between both datasets (pre-processed *vs* post-processed) using Levene’s test (Levene, 1960) where variance distributions were centred at their respective medians (H_0_:σ_(*a*)_^2^=σ_(*b*)_^2^;H_A_:σ_(*a*)_^2^≠σ_(*b*)_^2^). A Bonferroni-Dunn correction was applied (*n* = 16), and a *P*_BD_ ≤ 0.005 was considered statistically significant (Table 2). Levene’s test was chosen over more commonplace variance tests (such as an *F*-test) to account for potential deviations from a Gaussian distribution.

The each individual *P*_BD_ set (those derived from the Levene’s test and those derived from the Kruskal-Wallis test, respectively) were combined (*P*_F_) using the Fisher’s method (Fisher, 1938). As *P*_F_ was derived from *P*_BD_, a Bonferroni-Dunn correction was not applied in this instance and a *P*_F_ was considered statistically significant (Table 2). This process should be considered less important than the “contamination” reduction process.

#### Chosen metric

The median imputed dataset displayed the best performance for this analysis (Tables 1-2; Figures 3-4) and data derived from this impution metric were used for all subsequent analyses. Principal component analyses using raw data and processed data are presented for illustrative purposes (Figures 5-6).

**Figure 3:**
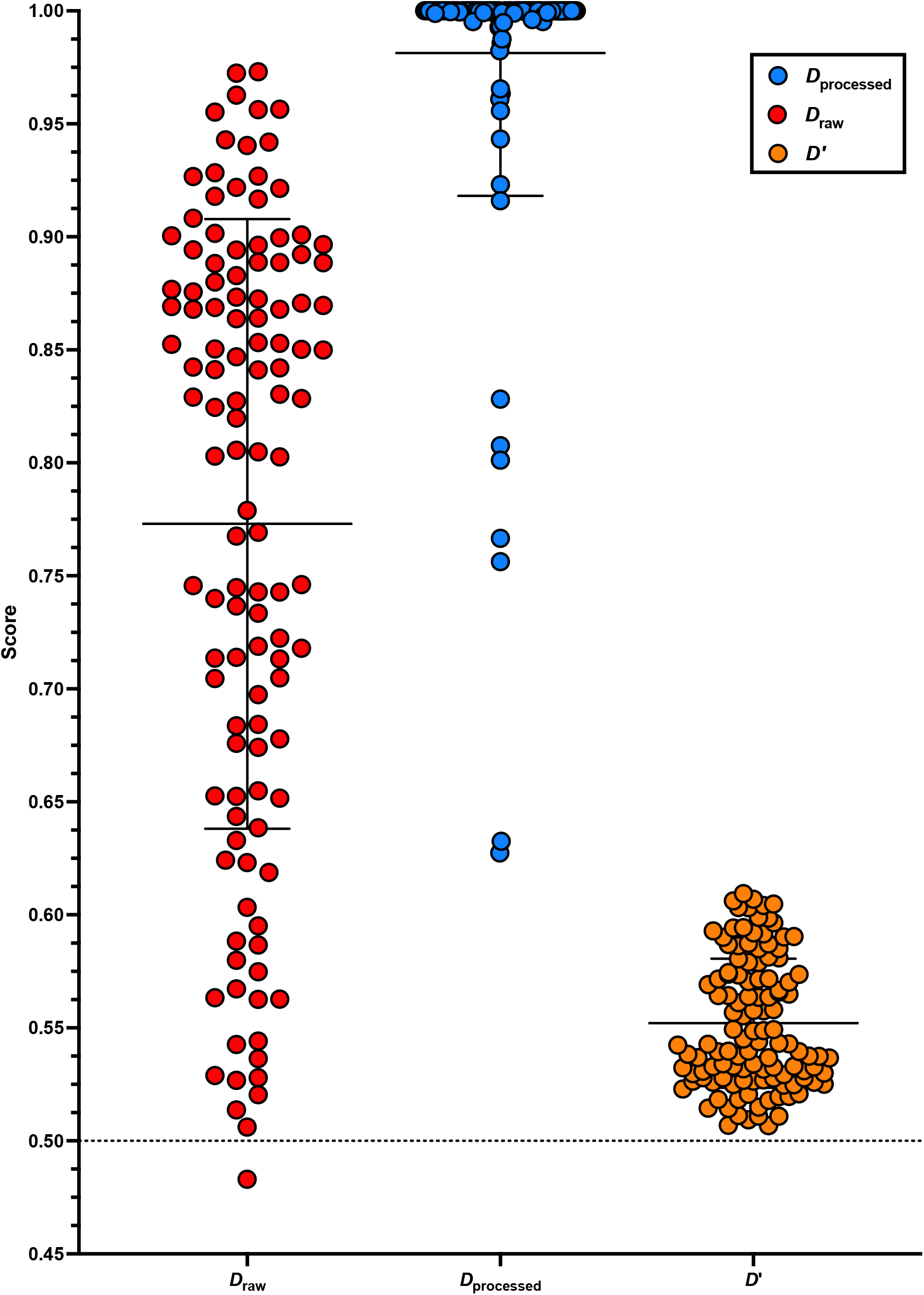
A comparison of distances. A comparison of distances between group pairs in the raw dataset (*D*_raw_) and processed dataset (*D*_processed_) with their associated performance scores (*D’*). The horizontal dashed line indicates the point at which an improvement between *D*_raw_ and *D*_processed_ is achieved. Long (full) horizontal lines behind each cluster indicate the mean and shorter lines indicate the standard deviation.

**Figure 4:**
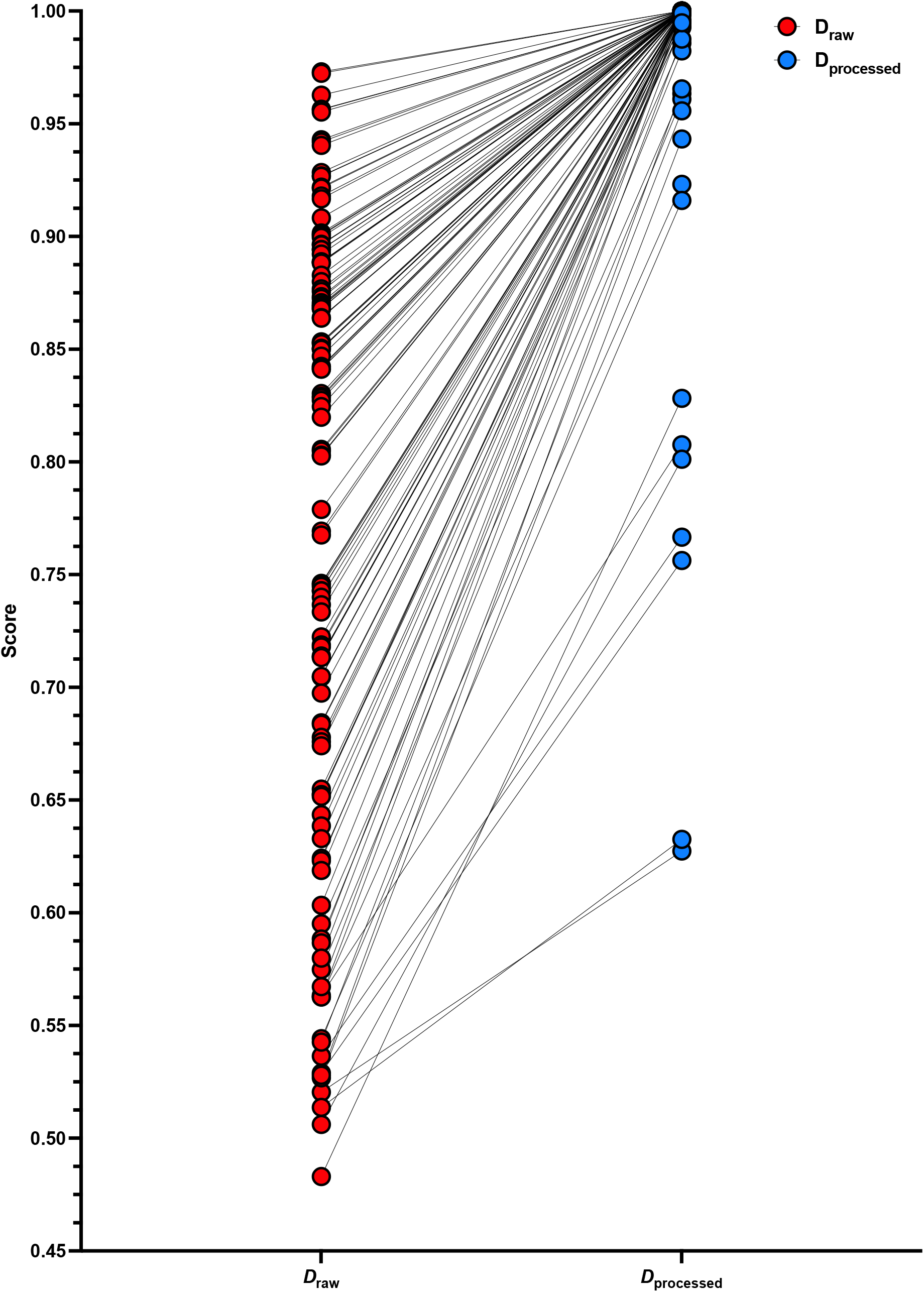
Paired comparison of distances between the raw and processed datasets. The effect of outlier processed is demonstrated with the line linking *D*_raw_ and *D*_processed_ for a given group pair. In every instance, *D*_processed_ is higher than its associated *D*_raw_ indicating greater separation.

**Figure 5:**
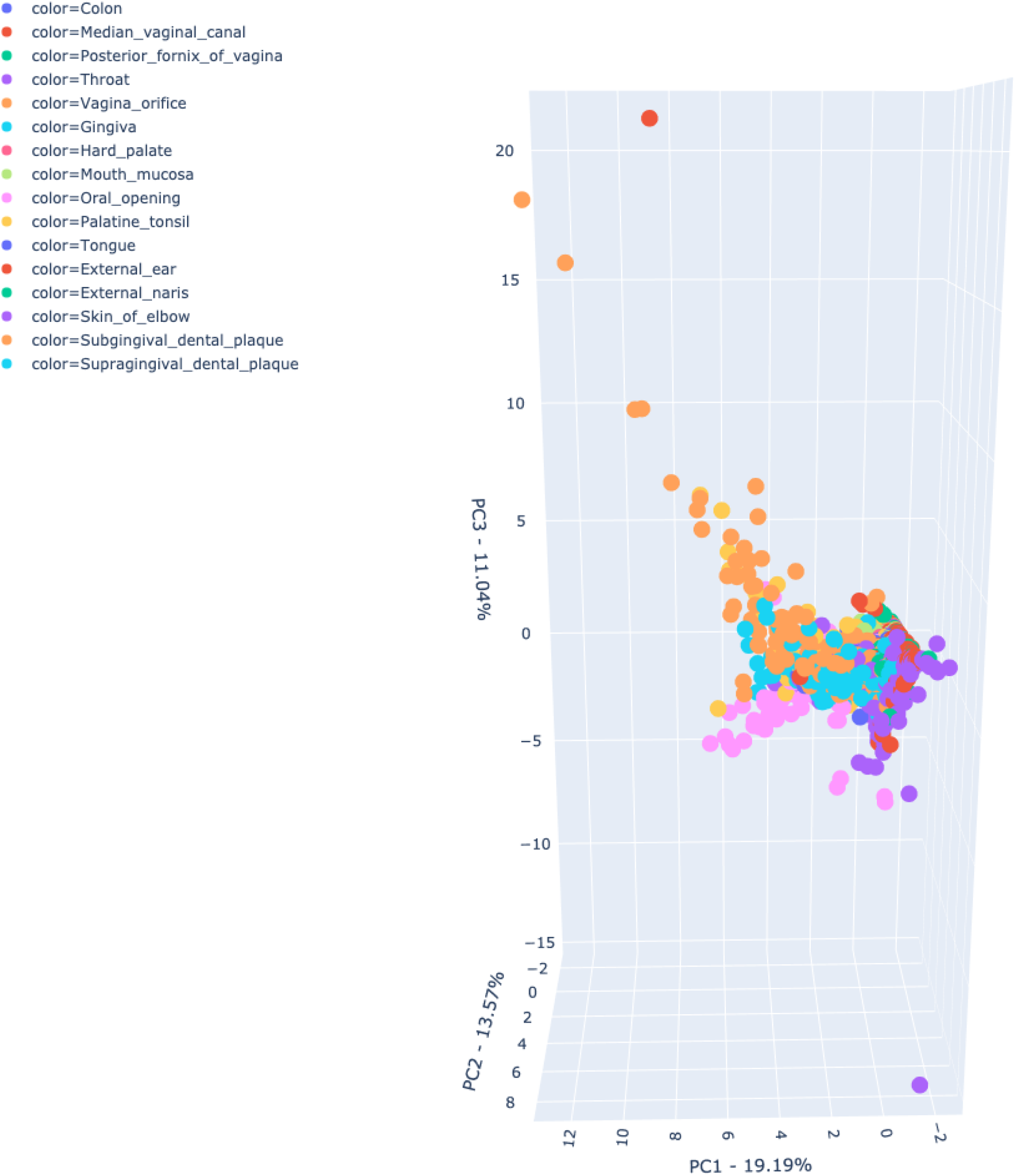
Principal component analysis of the raw (unprocessed) data. Sites are highly overlapping with considerable variance across the dataset and especially in the sebum category.

**Figure 6:**
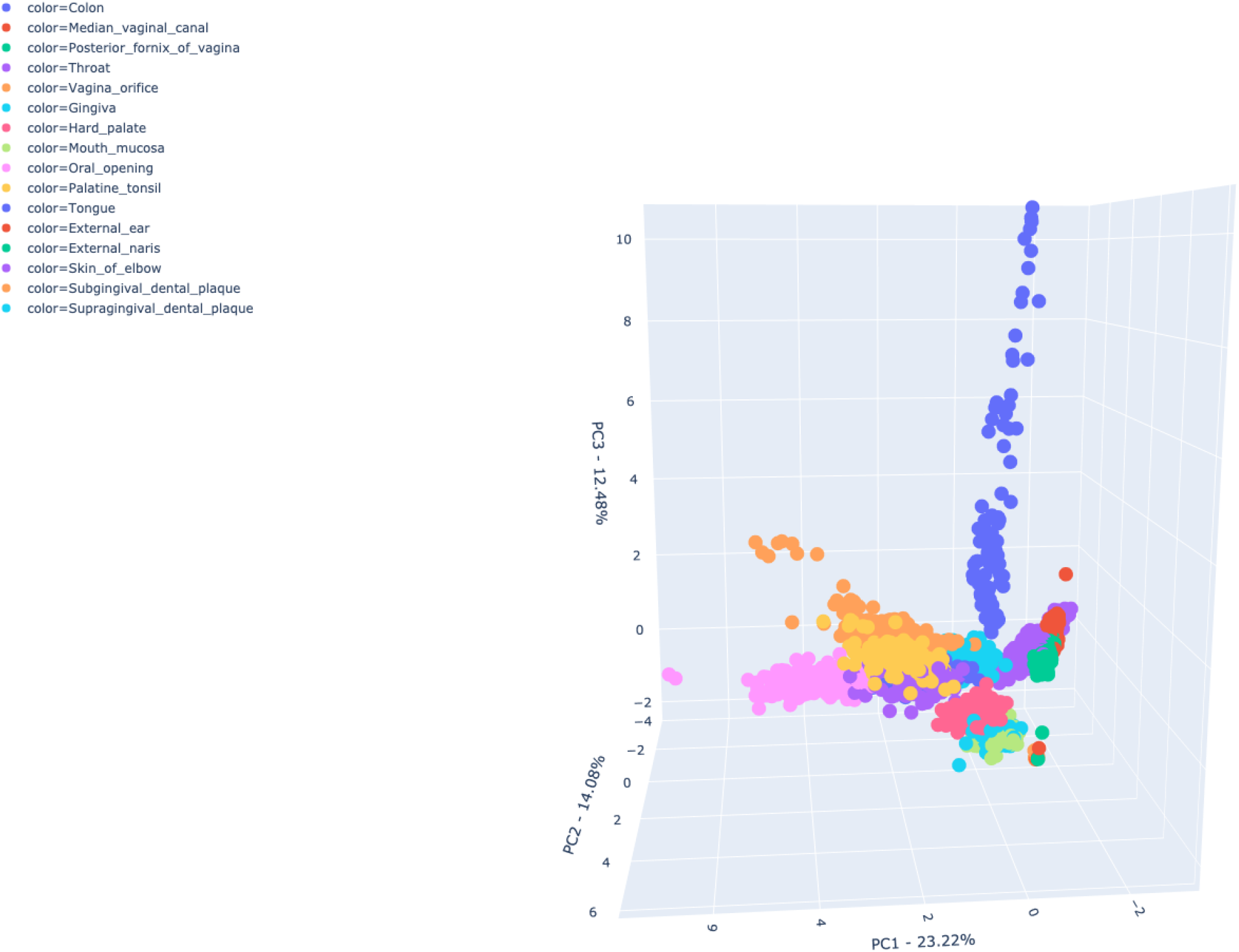
Principal component analysis of the processed data. Sites are much less overlapping with extensive overlap not occurring between many sites. There is still some variance across the data but is much less pronounced (as evidenced by the PC boundaries).

#### Statistical analysis

Descriptive statistics (mean, standard deviation, harmonic mean, geometric mean, median, variance, and standard error) were calculated for unprocessed and processed data (Table 3). For brevity, Phyla in a given group with a mean of 0 post-processing were removed from both groups (both pre- and post-processing). Treatment effect (comparison of variation) was assessed for each Phylum between both datasets (pre-processed *vs* post-processed) using Levene’s test where variance distributions were centred at their respective medians. A Bonferroni-Dunn correction was applied (*n* = 115) and a *P*_BD_ ≤ 0.005 was considered statistically significant (Table 3).

#### Reporting of changes made to the processed dataset

Following good practice, uniForest reports the descriptive statistics for the raw and processed datasets for each taxon in each group in addition to the proportion of imputed outliers (Table 3). For illustrative purposes microbiome data at Phylum level for the “Colon” group, one group from the skin (“Exterior nares”), one group from the mouth (“Palatine tonsil”) and one group from the vagina (“Median vaginal canal”) were randomly selected and their relative abundances (of both the pre- and postprocessed datasets) were displayed as an example (Figure 7). Relative abundances were made for any group where the proportion of outliers exceeded 0.7 (70%) to confirm the presence of multiple distinct subgroups within a given group (Figure 8).

**Figure 7:**
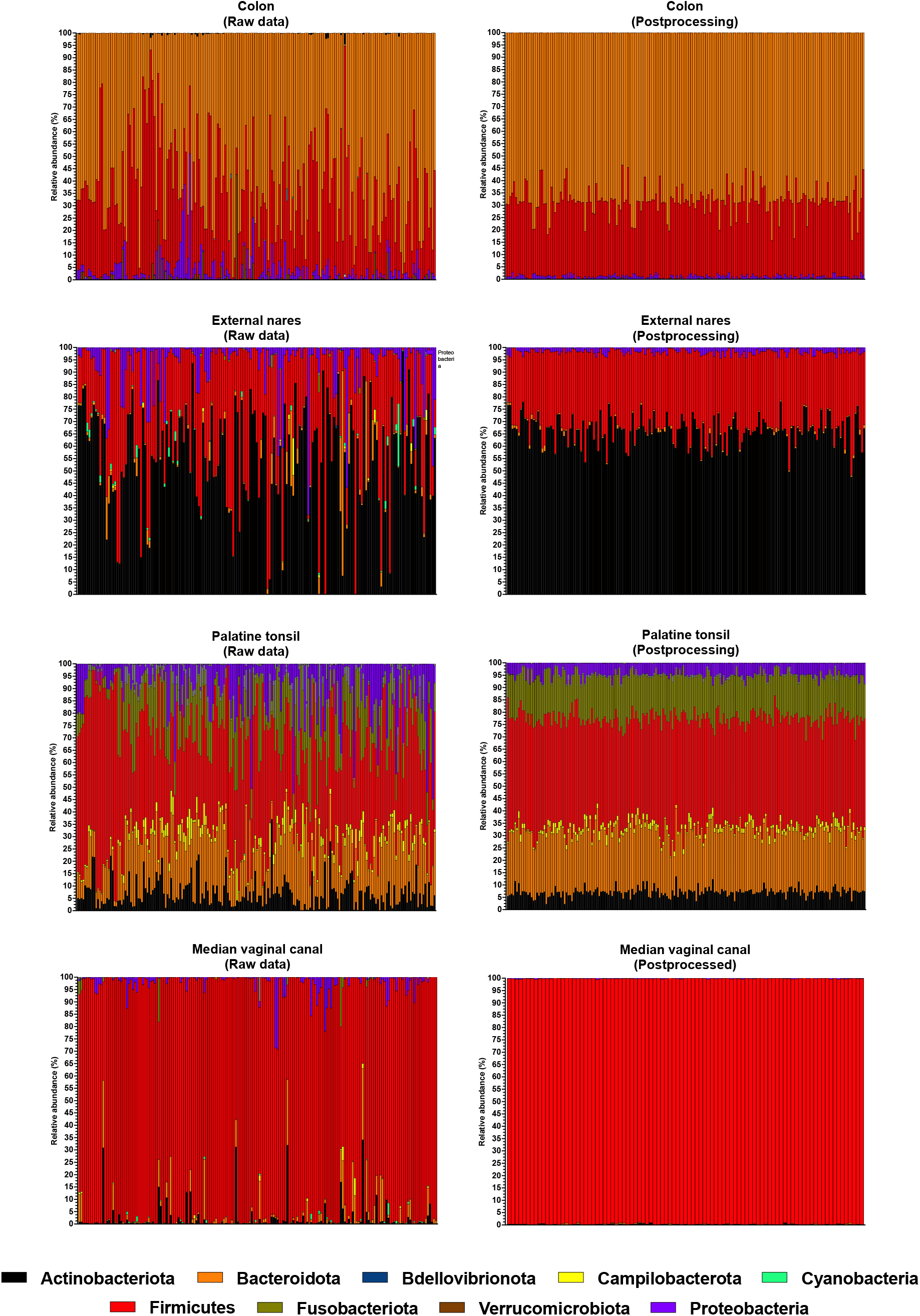
Effect of outlier impution on four anatomical groups. Samples (not numbered) are displayed along the *x*-axis and their relative abundances on the *y*-axis

**Figure 8:**
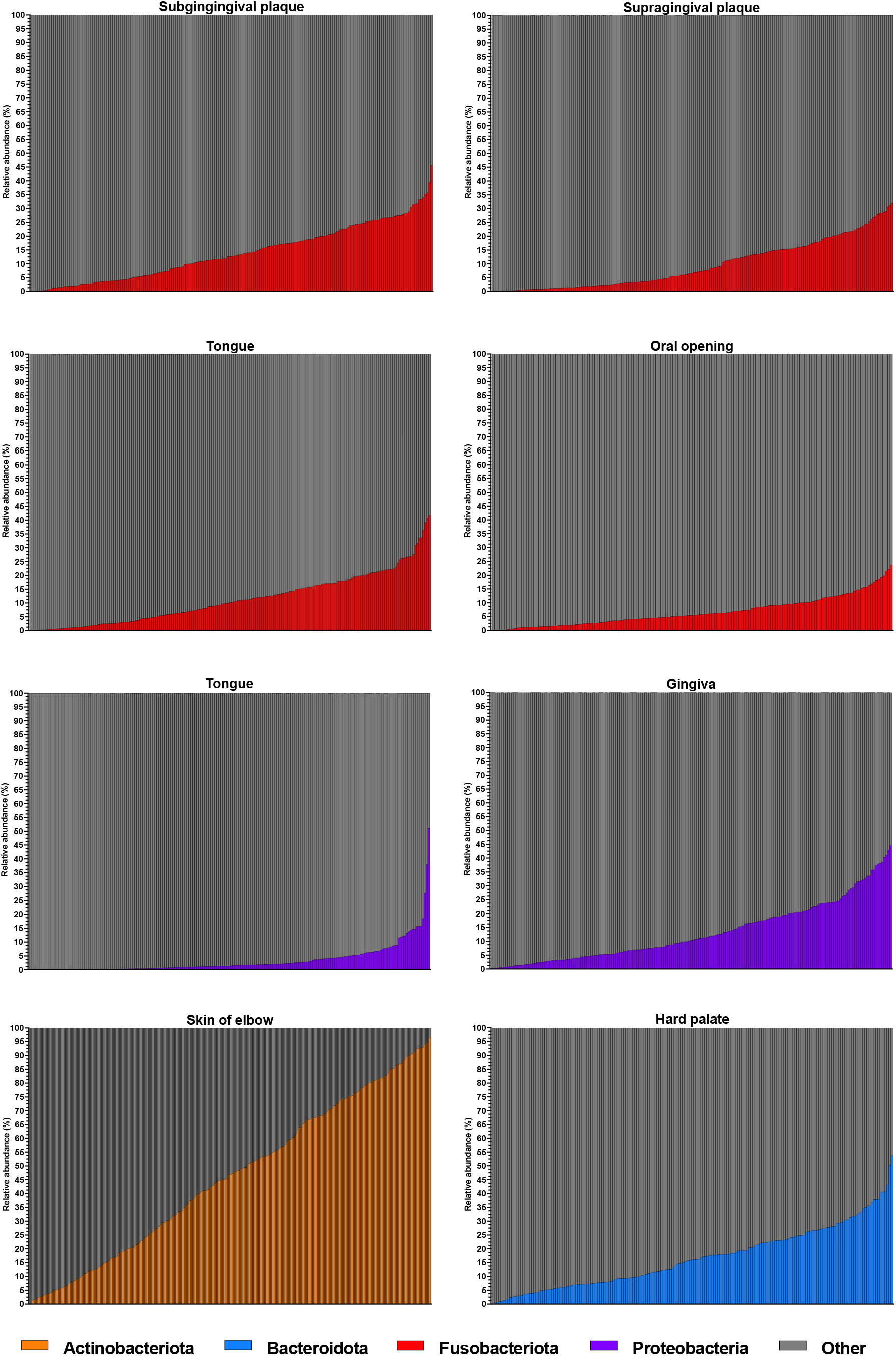
Distribution of Phyla in samples where impution exceeded 0.75 (75%). Samples (not numbered) are displayed along the *x*-axis and their relative abundances on the *y*-axis. Data is ordered from smallest to largest along the *x*-axis

#### Assessment of effect on α-diversity variation

A cohort of 4 α-diversity metrics (specifically Simpson’s *D*, Simpson’s *E*, Chao1, and Shannon’s *H* (Shannon, 1948; Simpson, 1949; Rényi, 1961; Chao, 1984)), were calculated for each sample in each group using the “math.diversity.alpha” function in the skbio Python library *v*.0.1.3 (http://scikit-bio.org/). Descriptive statistics (mean, standard deviation, median, variance, harmonic mean, geometric mean, and coefficient-of-variation) were calculated for both raw and processed datasets (Table 4; Figure 9). The difference (Δ) in coefficients-of-variation (CoV) were calculated using the formula:

**Figure 9:**
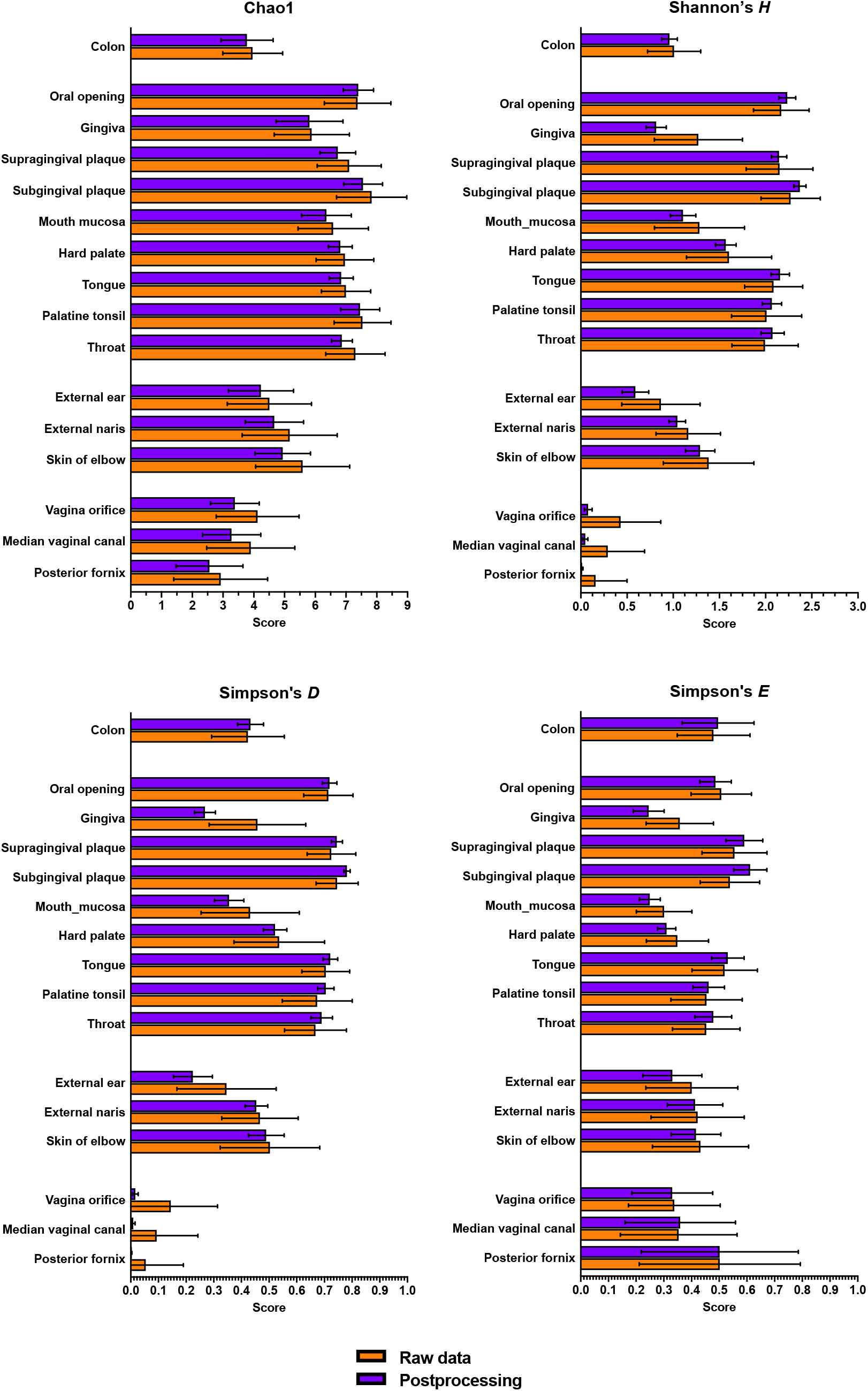
Comparison of *α*-diversity metrics.

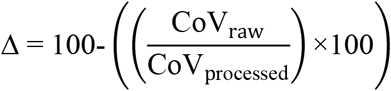

Statistical differences were determined using Levene’s test where variance distributions were centred at their respective medians. A Bonferroni-Dunn correction was applied (*n =* 64) and a *P* ≤ 0.005 was considered statistically significant (Table 4).

## Results

### Isolation Forests significantly reduces variance in reads across the dataset

Isolation Forest proved to be a powerful tool for reducing variance in microbiome datasets, achieving a statistically significant decrease (*P*_BD_ ≤ 0.005) in 87 of 115 (75.62%) comparisons (Table 3). A minimum reduction in variance of 81.16% was achieved. Examples of variance reduction are presented in Figure 6.

### Isolation Forests resolve groups with surprising accuracy

Datasets without outlier processing were highly overlapped irrespective of whether phyla were removed to match phyla removed by outlier processing (Figure 4). Following outlier removal, nodes within groups were tightly clustered together yet divergent from other groups. Data sampled from the vagina (“Median vaginal canal”, “Posterior fornix of vagina”, and “Vaginal orifice”) were highly overlapping, as were some samples from the buccal cavity (specifically the “Throat” and “Mucus membrane”). Relatively little overlap was observed beyond these though. When the distances between samples in a given group were compared, significant variation reduction (*P*_BD_ ≤ 3.46*e*^-82^) was observed in every comparison. This difference is not surprising when the data presented in Figures 4 and 5 are visually compared.

### Isolation Forests significantly reduces variance in α-diversity calculations

Isolation Forest significantly reduced (*P*_BD_ ≤ 0.005) variance in α-diversity in 55 of 64 (85.94%) of comparisons (Table 4), where Chao1 accounted for 4 of the 9 samples that were not statistically significant, Simpson’s *E* accounted for 4, and Simpson’s *D* accounted for 1). Simpson’s *E* was observed to be co-insignificant with Chao1 in 3 of 4 instances, and the instance where Simpson’s *D* was observed to be insignificant (‘Posterior fornix of vagina’) was also co-insignificant with Simpson’s *E*. Instances reported to have a significant reduction had lost between 17.68-85.69% in variance (Figure 6).

## Discussion

Isolation Forest has demonstrated considerable prowess in outlier processing across a multitude of fields (*eg*. Alonso-Sarria *et al*., 2019; de Santis & Costa, 2020; Elnour *et al*., 2020). To our knowledge, this is the first demonstration of its capabilities on microbiome datasets. In our opinion, Isolation Forests surpassed all expectations by significantly reducing variance across all Phyla in almost all instances (Table 2) and in reducing distance variation between samples in a given group (Table 3). While the variance reduction is most useful and may uncover otherwise overlooked trends, we were particularly surprised by its capacity to distinguish anatomical sites from others at the Phylum level. Interestingly, even as sites are grouped (*eg*. the external ear, external naris, and elbow are grouped as “sebum” by the original publication), excellent separation was achieved *via* PCA (Figure 2). Most interestingly, was the distinct separation of subgingival plaque and supragingival plaque as their geographical locations are mere millimetres apart. Both plaque groups displayed higher levels of variance (comparative to other groups) post-processing. This variance could be due to a myriad of overlapping reasons prior to sampling, such as how much time has passed since the subject last consumed food or liquid, the regular diet of the subject, how much time passed since the patient engaged in a dental hygiene activity, whether the subject used a biocidal mouthwash (*e*.*g*. zinc acetate), and the underlying dental anatomy of each subject, and the hygiene habits of each subject (Chen and Jiang, 2014; Suzuki *et al*., 2018; Burcham *et al*., 2020). As plaque is a biofilm produced by a cohort of bacterial communities, their underlying composition is likely more recalcitrant than other regions of the oral cavity so would be harder to control for experimental bias (Adler *et al*., 2013; Buckley *et al*., 2014; Velsko *et al*., 2019). Despite these variances however, Isolation Forest separated these sites with no overlap at the Phylum level. Using Isolation Forests for oral microbiome studies may aid in aetiological diagnostics in oral disease, especially for non-culturable organisms such as those in the *Candiatus* Patescibacteria Phylum.

In 33 of the 193 (33.85%) sample groups, more than 50% of the group dataset for a given taxon was detected as contamination, determined to be an outlier, and imputed (Table 3). Of the top 8 most imputed Phyla per group, 7 were sampled from the oral cavity. One particular Phylum (Fusobacteria) was subjected to impution over 70% of the time in 4 body sites, the ‘Tongue’, the ‘Oral opening’, the ‘Subgingival dental plaque’ and the ‘Supragingival dental plaque’. Proteobacteria were imputed over 80% of the time on the ‘Gingiva’, and ‘Tongue’. Bacteroidota was imputed over 70% of the time on the ‘Hard palate’. Finally, Actinobacteria were imputed over 80% of the time on the ‘Skin of elbow’. When the relative abundances of a given Phylum-of-interest was computed and arranged from smallest-to-largest (compared to all other Phyla in the group), a distinct pattern was observed: the data rose steadily with rare (if any) large spikes (Figure 5). This suggests that subgroups exist within the given groups, especially in parts of the oral cavity. Due to the sensitivity of iForests using the ‘auto’ contamination score, these inclines would have made the creation of a ‘normal space’ difficult. As 7 of the 8 most imputed samples arose in the mouth, with 3 groups (sites) recorded, multiple oral subgroups were suspected. Dysbiosis in oral Fusobacteriota, Bacteriota, and Proteobacteria are commonly associated with periodontitis, gingivitis, and periodontitis, and in the development of dental caries (Kapatral *et al*., 2002; Tanaka *et al*., 2008; Chen *et al*., 2017; Jiang *et al*., 2019). As these oral conditions vary in severity, underlying microbial dysbiotic variability can reasonably be expected. It is hypothesised that a cohort of subgroups exist within the oral population used in this study and preliminary separation of oral groups into subgroups prior to outlier processing is expected to reduce the rate of impution.

The rampant variance observed in the Actinobacteria relative abundances in the ‘Skin of elbow’ group could be due to a variety of factors such as hydration and sebum levels (Mukherjee *et al*., 2016; Lee *et al*., 2018) which could be easily augmented by factors such as clothing, weather, underlying condition, or age.

## Conclusion

Isolation Forests and simple impution of the mean have illustrated their potential prowess in microbiome studies. This procedure has proven useful in a human dataset and is expected to be useful for any studies where high levels of microbiome community variance is expected. It would be interesting to observe how this procedure affects ecological samples, such as for ecological validation.

## Code availability

The software and dataset used for this publication are fully available at https://github.com/RobLeighBioinformatics/MicrobiomeOutliers

